# Heat stress negatively affects betalain biosynthesis in *Amaranthus tricolor* through down-regulation of *CYP76AD1* and *5GT*

**DOI:** 10.1101/2025.03.05.641739

**Authors:** Ya-Ping Lin, Tomomi Ohokubo, Akito Nashiki, Yosuke Yoshioka, Yuan-Yun Zhang, Hieng-Ming Ting, Sachiko Isobe, Kenta Shirasawa, Ken Hoshikawa

## Abstract

Orphan crops are considered essential climatic-resilient genetic resources due to their native adaptability to various ecological environments. *Amaranthus tricolor* is one of a few common vegetables widely cultivated in tropical and subtropical regions. Strong vigor, abiotic-stress adaptability, and nutrition richness make it an ideal vegetable to provide food and nutrient security for smallholders. However, the breeding efforts depend on the field yield trials because little genomic information or genetic resources can be accessed. To fill this gap, we first used the Illumina and Pacbio sequencing systems to assemble a novel reference genome of *A. tricolor* using a Japanese variety Bayam. Together with a core collection developed in our previous study, we investigated the genetic mechanism of betalain biosynthesis under optimal and heat-stress conditions. Betalain is an antioxidant phytonutrient commonly appearing in beets and amaranths. As a result, ATR1.0CH03_5131103 which associate with *ATR1.0ch03g000565* was identified in the optimal condition. Meanwhile, according to the transcriptomic profiles in this study, *ATR1.0ch03g000565* also show significantly differential expressions under control and heat stress conditions, suggesting this candidate gene regulates the optimal performance of *CYP76AD1*. Meanwhile, among the homologous genes involving in the betalain biosynthesis, *CYP76AD1* and *5GT* were down-regulated under heat stress conditions; *cDOPA* was up-regulated. This result suggested heat stress reduces betacyanin biosynthesis through the downregulation of *CYP76AD1* and *5GT*; despite that the upregulation of *cDOPA* cannot rescue the decrease of betalain. More molecular experiments are required to address the regulation of these three enzymes.

## Introduction

Malnutrition has long been a critical threat to human health, with an estimated 821 million people in South Asia and Africa suffering from inadequate food consumption and nutritional deficiencies (FAO, 2018). Orphan crops, which include various indigenous or local vegetables in developing countries, are increasingly recognized as valuable resources for establishing sustainable food systems that can provide nutritious foods in an environmentally resilient manner (Sarker et al., 2022). One of the indigenous vegetables, *Amaranthus*, has been revealed to be able to well-adapted to a range of abiotic stresses, including drought, salinity, heat, and heavy metal exposure (Sarker et al., 2022; Reyes-Rosales et al., 2023). The genus *Amaranthus* comprises approximately 50‒70 species globally, some of which are consumed as pseudo-grains, others as leafy vegetables (Sarker et al., 2022). *Amaranthus tricolor* has traditionally been cultivated in Africa and Asia, where they are valued as low-input, nutrient-rich, and medicinal vegetables (Sarker et al., 2022). It is often used as a substitute for spinach due to its high content of carotenoids, vitamin C, minerals, and flavonoids (Sarker et al., 2022). Their high nutrient content and tolerance to abiotic stresses make leafy amaranths promising candidates for climate-resilient agriculture.

Most breeding efforts in *A. tricolor* have focused on improving yield, with breeders selecting relatively stable and high-yield lines based on field trials. However, advances in high-throughput sequencing technologies have accelerated the study of functional genes in *A. tricolor*. For instance, transcriptomic analyses have identified *AmCYP76AD1* as a crucial gene in betalain biosynthesis, providing a pathway for improving betalain content through marker-assisted selection or further functional studies (Chang et al., 2021). A recent study used selected-genome sequencing approach to develop single-nucleotide polymorphisms (SNPs) for a worldwide *A. tricolor* collection and to develop a core collection (Hoshikawa et al., 2023). However, due to the absence of a reference genome for *A. tricolor*, SNP marker development has relied either on the reference genome of *A. hypochondriacus* or on *de novo* SNP calling. Given that *A. tricolor* is genetically distant from *A. hypochondriacus* and possesses a different chromosome number, mapping rates to the *A. hypochondriacus* reference genome are relatively low (Nguyen et al., 2019; Lin et al., 2021), consequently reducing the SNP number.

Betalains are unique pigments found in most species within the order *Caryophyllales*, including the family *Amaranthaceae* (Sarker et al., 2022). As phytonutrients, betalains have pharmaceutical applications, with intermediate products in the betalain pathway being used in the treatment of Parkinson’s disease (Galanie et al., 2015). Additionally, betalains are natural colorants widely used in the food industry (Galanie et al., 2015). In plants, betalains have diverse functions, including attracting pollinators, repelling herbivores, protecting tissues from ultraviolet radiation, and removing reactive oxygen species (ROS), all of which contribute to abiotic stress tolerance (Albano et al., 2015). Betalains are classified as betacyanins and betaxanthins, which are red and yellow, respectively. Betacyanins include four types of pigments, namely, amaranthin, betanin, gomphrenin, and decarboxybetanin (Sarker et al., 2022). L-tylosin, a precursor of betalain biosynthesis, is supplied by the shikimate pathway (Liu et al., 2019). The betalain biosynthesis pathway involves five key enzymes, including cytochrome P450 76AD1 (*CYP76AD1*: *LOC130803140*), L-dihydroxyphenylalanine 4,5-dioxygenase (*DODA*: *LOC130803136*), cyclo-DOPA 5-O-glucosyltransferase (*cDOPA*: *LOC130809389*), betanidin-5-O-glucosyltransferase (*5GT*: *LOC130825231*), and tyrosine decarboxylase (*TyDC*: *LOC130817294*) (Chang et al., 2021; Figure 1). *CYP76AD1* serves as the first enzyme to initiate the betalain biosynthesis, which is followed by either through *DODA* and *5GT* or *CYP76AD1* and *cDOPA* to produce betanin, and via *DODA* to synthesize betaxanthin.

**Figure 1.**
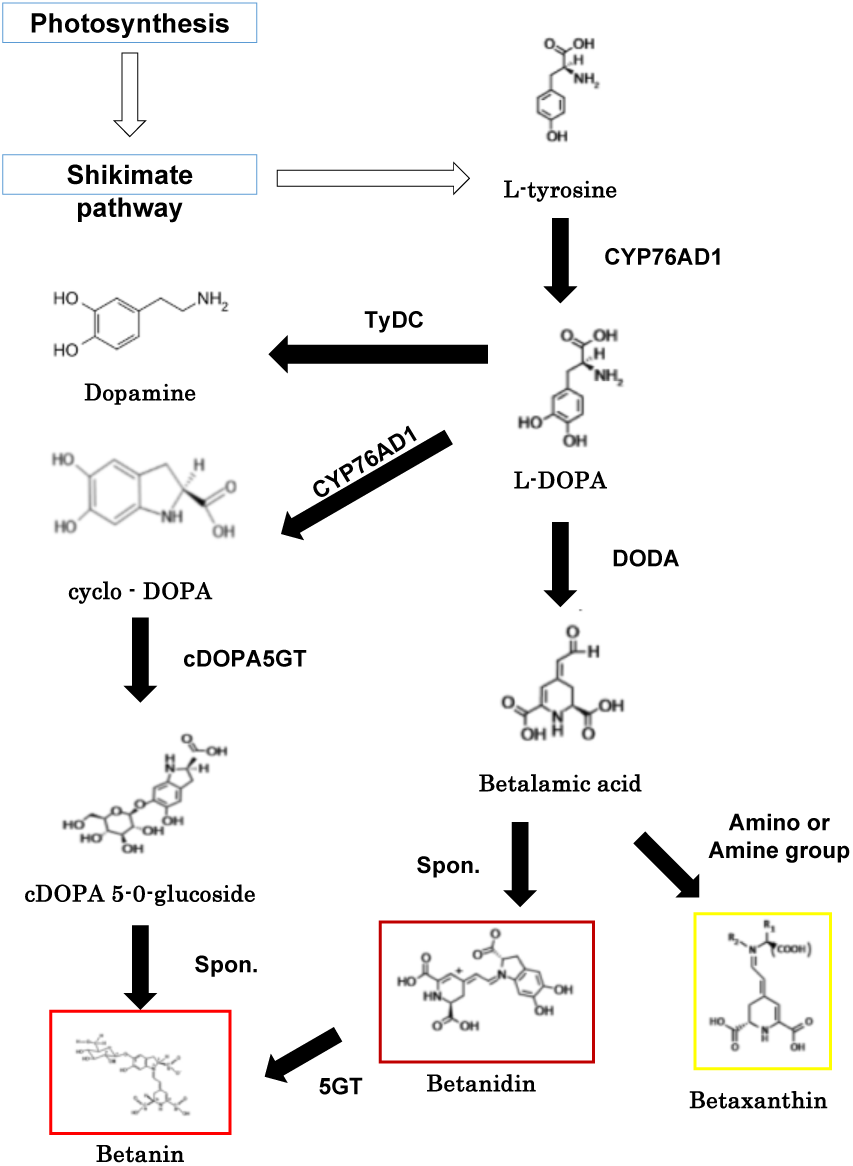
Pathway of betalain biosynthesis. Five enzymes participate in the betalain biosynthesis are as following: *CYP76AD1* (*cytochrome P450 76AD1*), L-DOPA (L-3,4-dihydroxyphenylalanine), *DODA* (*L-dihydroxyphenylalanine 4,5-dioxygenase*), *cDOPA5GT* (*cyclo-DOPA 5-O-glucosyltransferase*), *B5GT*, (*betanidin-5-O-glucosyltransferase*), and *TyDC* (*tyrosine decarboxylase*).

Additional transcription factors are involved in the regulation of betalain biosynthesis. Previous studies have highlighted the WRKY family members, which were not only abundant in the amaranth transcriptomic profile but also demonstrated the ability to bind to the *CYP76AD1* promoter, thereby promoting betacyanin synthesis (Yang et al., 2024). Overexpression of *AtrWRKY42-2* has been shown to upregulate *CYP76AD1*, *B5-GT*, and *B6-GT*, and *TyDC* (Yang et al., 2024). Similar findings were reported in pitaya (*Hylocereus monacanthus*), underscoring the role of WRKY transcription factors in betacyanin biosynthesis (Zhang et al., 2020). Moreover, the MYB family has also been implicated in the regulation of betacyanin biosynthesis. In pitaya, *HuMYB132* enhances the transcriptional activity of *HuCYP76AD1–1* and *HuDODA1* by binding to their promoters (Xie et al., 2023). Silencing *HuMYB132* leads to reduced betalain accumulation and downregulation of betalain biosynthetic genes. In pear, *PyWRKY26* has been found to target *PyMYB114*, thereby regulating anthocyanin accumulation (Li et al., 2020). These findings suggest a complex, yet relatively unexplored, regulatory network governing betalain biosynthesis.

The objectives of this study are to investigate whether the betalain content of *A. tricolor* varies across different cultivation seasons and to elucidate the genetic mechanisms underlying betalain biosynthesis under heat stress conditions. However, due to the limited genomic information available for *A. tricolor*, a comprehensive examination of the genetic mechanisms involved in betalain biosynthesis has been challenging. To address this, we assembled a novel reference genome using an elite Japanese cultivar, ’Bayam.’ Building on our previous research, we re-performed reference genome-based SNP calling to develop a new SNP dataset for the core collection. We conducted a genome-wide association study (GWAS) to identify quantitative trait loci (QTL) associated with betalain biosynthesis across winter and summer seasons. Additionally, transcriptomic profiles were analyzed to 1) refine GWAS candidate genes, 2) assess the expression of homologous genes of *5GT*, *cDOPA*, *CYP76AD1*, *DODA*, and *TyDC*, under both control and heat stress conditions, and 3) identify new candidate genes involved in the regulation of betalain biosynthesis.

## Materials and Methods

### Plant Material

This study utilized the core collection of *Amaranthus tricolor* established in a previous investigation (Hoshikawa et al., 2023; Supplementary Table 1) to examine betalain content across winter and summer seasons in Taiwan. A total of 95 lines, categorized into 60 green, 19 red, and 15 mixed-color variants, were cultivated in a greenhouse for phenotypic assessment. The winter cultivation took place in October 2021 in Tainan, Taiwan, where the recorded daily maximum, minimum, and mean temperatures were 31.2°C, 23.1°C, and 26.7°C, respectively. The summer cultivation occurred in August 2022, also in Tainan, with corresponding daily maximum, minimum, and mean temperatures of 33.8°C, 25.1°C, and 28.5°C. Plants were grown for one month, after which seedlings were harvested for betalain content analysis.

### Measurement of Betalain Content

Betalain content was quantified following the protocol outlined by Ravichandran et al. (2013). Samples were initially flash-frozen in liquid nitrogen and subsequently freeze-dried. For extraction, 0.1 g of freeze-dried material was dissolved in 10 ml of 50% ethanol. The mixture was then agitated and centrifuged at 6000 rpm for 10 minutes. The supernatant was collected, and the centrifugation process was repeated twice. Betalain content in the extracted solution was determined using a UV-VIS spectrophotometer at 538 nm (for betacyanins) and 480 nm (for betaxanthins). The total betalain content was calculated by summing the absorbance values at 538 nm and 480 nm. Each line was evaluated in triplicate (three biological replicates). To assess the seasonal variation in betalain content, an ANOVA test was conducted, treating season and plant color as fixed effects.

### Genome assembly of *Amaranthus tricolor* cv. Bayam

The selfing F_7_ generation of *A. tricolor* cv. Bayam was used as the model cultivar for *A. tricolor* to construct a novel reference genome. Illumina and PacBio sequencing systems were used to assemble a new chromosome-level reference genome. For Illumina sequencing, genomic DNA was extracted from young *A. tricolor* cv. Bayam leaves provided by NOGUCHI Seed Co. LTD, with a Genomic-Tip (Qiagen, Hilden, Germany). A short-read sequencing library was constructed using the TruSeq DNA polymerase chain reaction-Free LT Sample Prep Kit (Illumina, San Diego, CA, USA) and sequenced using the Illumina HiSeqX platform. K-mer frequency was used to estimate the genome size using jellyfish (Marçais and Kingsford, 2011). For PacBio sequencing, DNA libraries were constructed using the SMRTbell Express Template Prep Kit 2.0 (PacBio, Menlo Park, CA, USA) and sequenced for six Single-molecule real-time (SMRT) cells (1M v3) on the PacBio platform. The reads were assembled with Falcon Unzip (https://github.com/PacificBiosciences/FALCON/) to obtain primary contigs and haplotigs of the genome. Potential sequence errors in the assembled sequences were corrected twice using long reads with Arrow implemented in the SMRT Link (PacBio). The pseudomolecular sequences were constructed based on a genetic map developed from the F_2_ population (n = 380). The F_2_ population was derived from a cross between Bayam and Asian Red, and SNP calling followed the procedures described by Shirasawa et al. (2016). Briefly, genomic DNA was extracted from the leaves and digested with two restriction enzymes, *Pst*I and *Msp*I. Libraries were sequenced on a DNBSEQ-G400 platform (MGI). The sequencing reads were mapped onto primary contigs using Bowtie2 (Langmead and Salzberg, 2012). High-confidence biallelic SNPs were identified using the mpileup option of SAMtools (Li et al., 2009) and filtered using VCFtools (Danecek et al., 2011) under the following conditions: read depth ≥ 5, SNP quality = 10, minor allele frequency ≥ 0.2, and proportion of missing data < 50%. A genetic map was constructed using Lep-Map3 (Rastas, 2017). The contig sequences were anchored to a genetic map to build the pseudomolecular sequence ALLMAPS (Tang et al., 2015). Repetitive sequences were identified using the RepeatMasker (Smit et al.). In accordance with RepeatMasker, repeat elements were classified into nine types: short interspersed nuclear elements, long interspersed nuclear elements, long terminal repeat elements, DNA elements, small RNA, satellites, simple repeats, low-complexity repeats, and unclassified elements.

### Genome Annotation

To comprehensively investigate gene distribution across the entire genome, we performed a thorough annotation, encompassing both coding and repetitive regions. This annotation was based on a combination of ab initio predictions, RNA, and protein evidence. For ab initio predictions, gene models were generated using AUGUSTUS v3.4.0 with the “Arabidopsis” parameter (Stanke et al., 2008). RNA evidence was derived from transcriptomic data obtained from *Amaranthus* species grown under controlled conditions. These RNA-Seq reads were aligned to the newly assembled reference genome using HISAT2 v2.2.1 (Kim et al., 2019), and transcripts were subsequently reconstructed with StringTie v2.2.2 (Pertea et al., 2015). Open Reading Frames (ORFs) longer than 100 amino acids were predicted using TransDecoder v5.7.0, referencing UniProt and Pfam domains (Haas et al., 2017). Protein evidence involved aligning sequences from *Amaranthus hypochondriacus*, *Amaranthus cruentus*, and *Spinacia oleracea* to the new reference genome using GenomeThreader v1.7.1 (Gremme et al., 2005). The ab initio predictions, RNA evidence, and protein evidence were integrated with respective weights of 1, 5, and 5 using EVidenceModeler v1.1.1 (Haas et al., 2008) to construct consensus gene models. Functional annotation of the predicted genes was performed using eggNOG-mapper v2.1.9 (Cantalapiedra et al., 2021). Additionally, homologous genes related to betalain biosynthesis, including *CYP76AD1* (*QOP57914.1*), *DODA* (*QOP57916.1*), *cDOPA* (*QOP57917.1*), *5GT* (*AJY59053.1*), and *TyDC* (*AJW81117.1*), were identified through BLAST with an *E-value* threshold of < 10^-50.

### Genome-Wide Association Study (GWAS) of Betalain Content

We conducted a genome-wide association study (GWAS) to identify genetic variants linked to betalain content. Illumina reads from double digest restriction-site associated DNA sequencing (ddRAD-seq) were retrieved from the DDBJ Sequence Read Archive database (accession numbers DRA012903 - DRA012912). The reads were quality-filtered using SolexaQA v3.1.7.1 (Cox et al., 2010) and aligned to the newly constructed reference genome using BWA v0.7.17 (Li and Durbin, 2009). Single nucleotide polymorphisms (SNPs) were called using GATK v4.2.3.0 (McKenna et al., 2010) and filtered based on a minor allele frequency threshold of < 0.05. Prior to GWAS analysis, leaf color phenotypes were encoded as dummy variables. Two models, Blink and FarmCPU, were employed within the Genome Association and Prediction Integrated Tool (GAPIT) (Wang and Zhang, 2021) to identify quantitative trait loci (QTL) with a significance threshold set at *p-value* < 0.001.

### RNA Experiments

Seeds of *Amaranthus* accessions Bayam, Am087, and Am127 were sown in plastic trays and grown in a controlled growth chamber at 25°C with a photoperiod of 16 hours light and 8 hours dark. After one week, seedlings were subjected to either heat-stress conditions (35/25 °C, 16 h/8 h light/dark) or maintained under control conditions (25 °C, 16 h/8 h light/dark) for one month. Leaves from one-month-old seedlings were harvested for RNA sequencing, with three biological replicates per accession under each treatment. Each biological replicate comprised leaves from three individual plants. RNA extraction and sequencing were performed by Rhelixa (Tokyo, Japan).

### Identification of Differentially Expressed Genes (DEGs)

RNA sequencing reads were processed to remove low-quality sequences using Trimmomatic v0.39 (Bolger et al., 2014) and then aligned to the new reference genome using HISAT2 v2.2.1 (Kim et al., 2019). Differentially expressed genes (DEGs) were identified using the DESeq2 package in R (Love et al., 2014), with significance defined as an adjusted p-value < 0.05. Log2 fold changes in gene expression between heat-stress and control conditions were calculated. Gene Ontology (GO) enrichment analysis was conducted using TBtools-II (Chen et al., 2023), applying a Benjamini-Hochberg adjusted p-value threshold of < 0.05.

## Results

### Betalain Content Across Seasons

The betalain content in the core collection exhibited significant seasonal variation. During winter, the betalain content ranged from 66.95 to 305.10, while in summer, it ranged from 14.51 to 51.36, highlighting a general decrease in betalain biosynthesis under heat stress conditions (Figure 2, Supplementary Table 1). The Pearson correlation coefficient between betalain levels in the two seasons was 0.29 (*p-value* = 0.006), suggesting a moderate but statistically significant correlation. Notably, red amaranths demonstrated higher betacyanin levels in winter compared to green amaranths, while both types showed the lowest betacyanin concentrations in summer (Supplementary Figure 3). In contrast, betaxanthin levels were primarily influenced by seasonal changes (Supplementary Figure 3), indicating that red and green amaranths may employ distinct regulatory mechanisms for betacyanin and betaxanthin synthesis in response to elevated temperatures.

**Figure 2.**
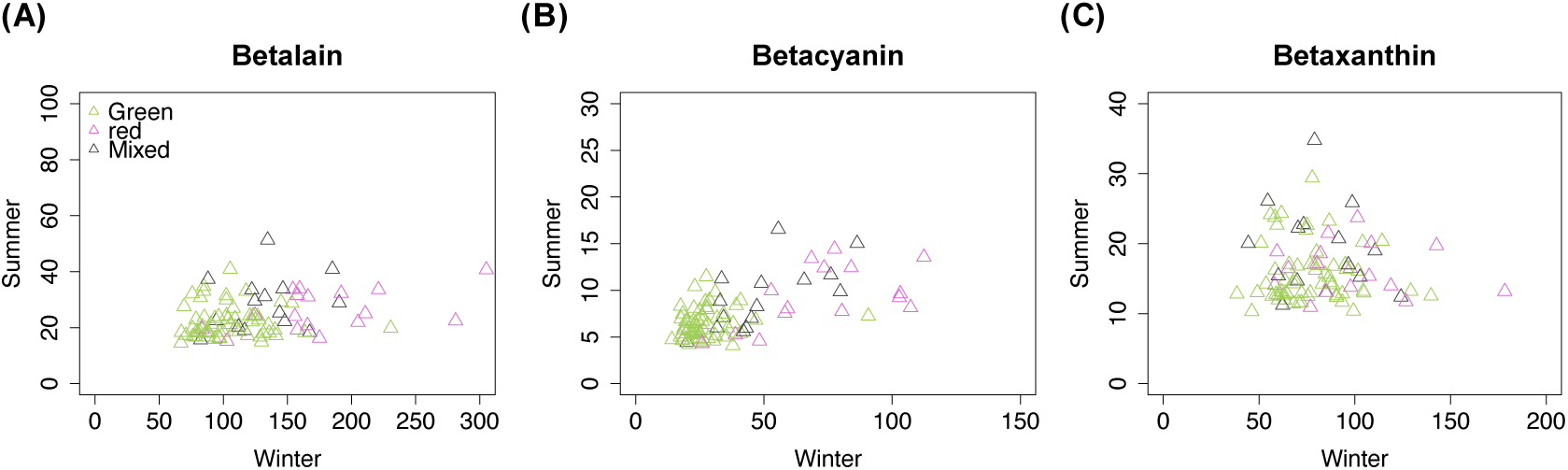
Correlation of betalain in the winter and summer seasons. (A), (B), and (C) are the total betalain, betacyanin, and betaxanthin, respectively. Green, violet, and grey indicate green, red, and mixed-color amaranths, respectively.

### Reference Genome of *A. tricolor* cv. Bayam

The reference genome for *A. tricolor* cv. Bayam was constructed using a combination of 126.7 Gb high-quality Illumina short reads and 57.6 Gb PacBio Continuous Long Reads. Genome size estimation, based on K-mer distribution analysis, was approximately 596.4 Mb (Supplementary Figure 2). PacBio sequencing yielded 57.6 Gb of reads with an N50 length of 38.1 kb, which were assembled into 2,151 contigs (Supplementary Table 2). A genetic map anchored 199 contigs to 17 chromosome-level scaffolds, spanning a total of 231.4 Mb (Supplementary Table 3). Repetitive sequences accounted for 52.9% of the genome, with LTR retroelements being the most dominant repeat type (8.9%), followed by simple repeats (3.9%) and DNA transposons (3.6%). In addition, a comprehensive sequence similarity search across the whole genome identified 5,599 pairwise homologous genes, forming 148 collinear blocks across 68 chromosome pairs (Supplementary Figure 3). Six pairs of these blocks were intra-chromosomal on chromosomes 1, 2, and 4, while the remaining 62 pairs were inter-chromosomal. Chromosome 2 exhibited 71 collinear blocks with all other chromosomes except for chromosome 16, whereas chromosome 14 shared only two homologous blocks with chromosome 2.

A total of 58,844 genes were annotated across the genome, including five genes encoding enzymes involved in betalain biosynthesis: *5GT*, *cDOPA*, *CYP76AD1*, *DODA*, and *TyDC*. These genes were blasted against the new reference genome to identify their homologous counterparts in *A. tricolor* (Supplementary Table 4). This search revealed 14, 18, 21, 2, and 5 homologous genes for *5GT*, *cDOPA*, *CYP76AD1*, *DODA*, and *TyDC*, respectively. Interestingly, *5GT* and *cDOPA* shared 12 homologous genes due to a 30% similarity in protein sequence (*E-value* = 10^-64). These homologous genes were assigned to either *5GT* or *cDOPA* based on the *E-value*, resulting in 12 and 8 *5GT* or *cDOPA* homologous genes, respectively.

### Candidate QTLs Regulating Betalain Biosynthesis

In this core collection, a total of 40K SNPs were initially identified, with 2,912 SNPs retained following filtering based on minor allele frequency. Principal Coordinate Analysis (PCA) demonstrated an absence of clear genetic clustering within this collection, suggesting a reduced likelihood of false-positive associations (Supplementary Figure 4). GWAS identified 39 and 6 significant SNPs associated with betalain content in the winter and summer seasons, respectively (Figure 3, Supplementary Figure 5, Supplementary Table 5). Of these, 12 SNPs, seven in winter and five in summer, were consistently detected by both the FarmCPU and Blink models, thereby strengthening confidence in these loci (Table 1). The SNP ATR1.0CH17_6168377 was identified in association with both betacyanin and betaxanthin during the winter season, while ATR1.0CH00_181109920 and ATR1.0CH12_6719988 were detected in association with both pigments during the summer season. These findings suggest that betalain biosynthesis is regulated by distinct mechanisms under varying seasonal conditions, with a significant genotype-by-environment interaction effect. Candidate genes were defined as those either directly targeted by these significant SNPs or located proximally upstream or downstream (Supplementary Table 6). Notably, none of the 115 candidate genes identified have been previously implicated in betalain biosynthesis, indicating the need for further investigation into their potential roles.

**Figure 3.**
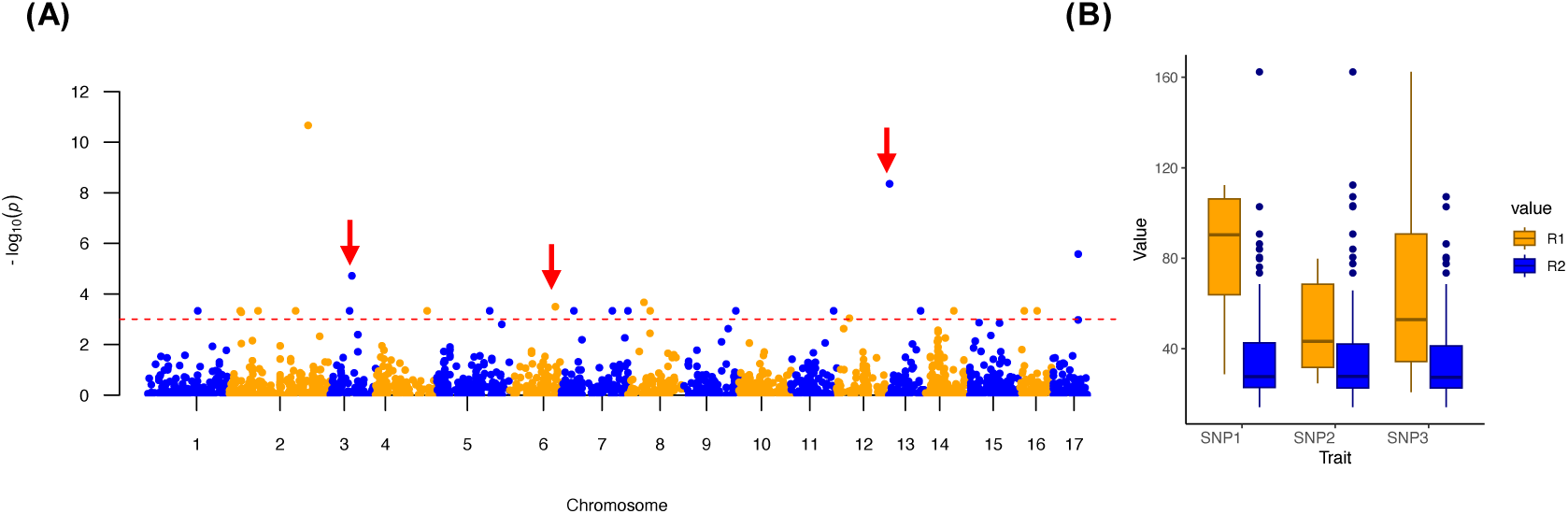
GWAS of betacyanin content and the impact of candidate SNPs. (A) The Manhattan plot for betacyanin levels during the optimal season, as analyzed using the FarmCPU model. Red arrows indicate significant SNPs associated with DEG s. (B) The effects of SNPs linked to the DEGs. The SNPs denoted as SNP1, SNP2, and SNP3 correspond to ATR1.0CH03_5131103, ATR1.0CH06_11090793, and ATR1.0CH13_174259, respectively.

**Table 1.**
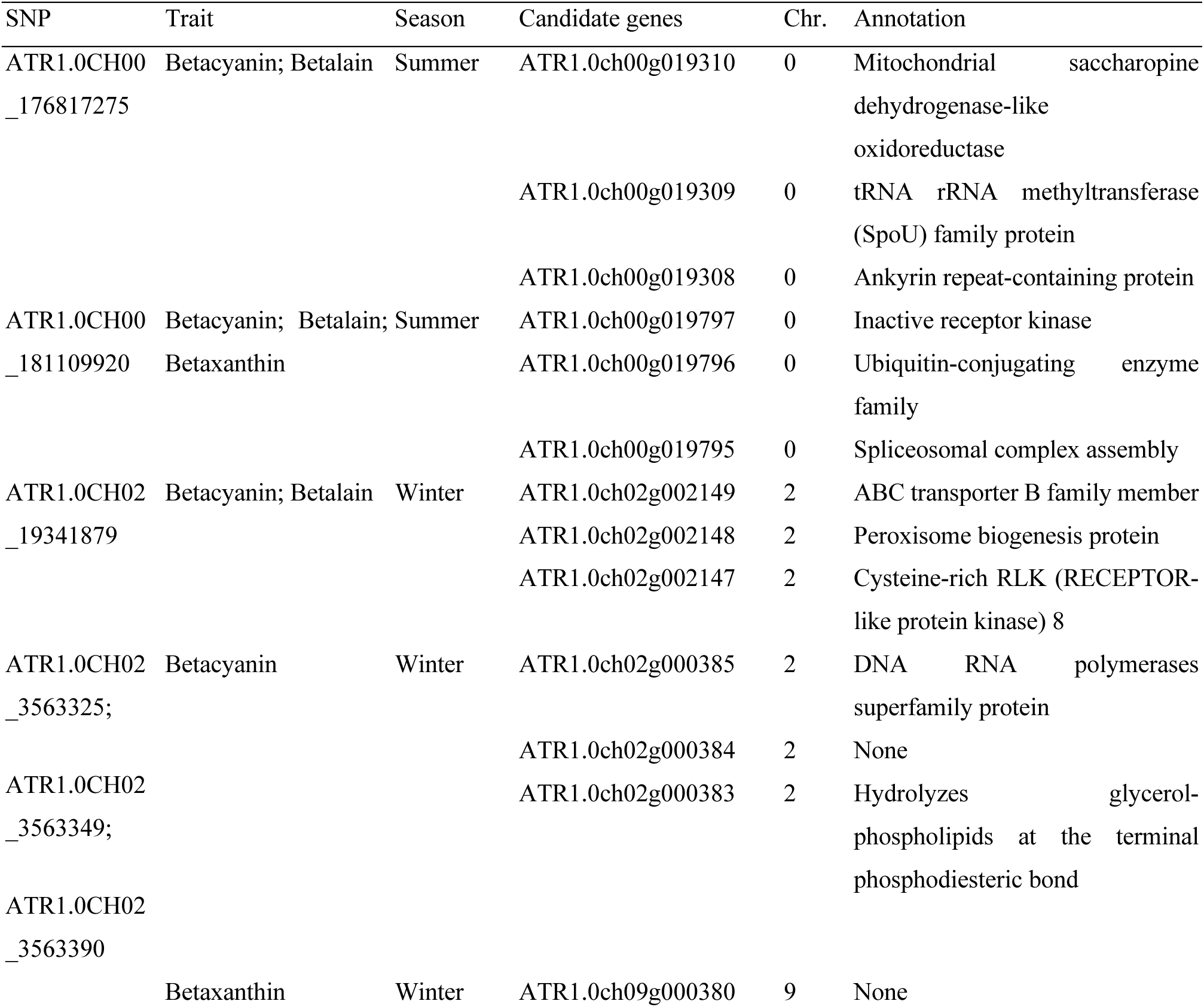
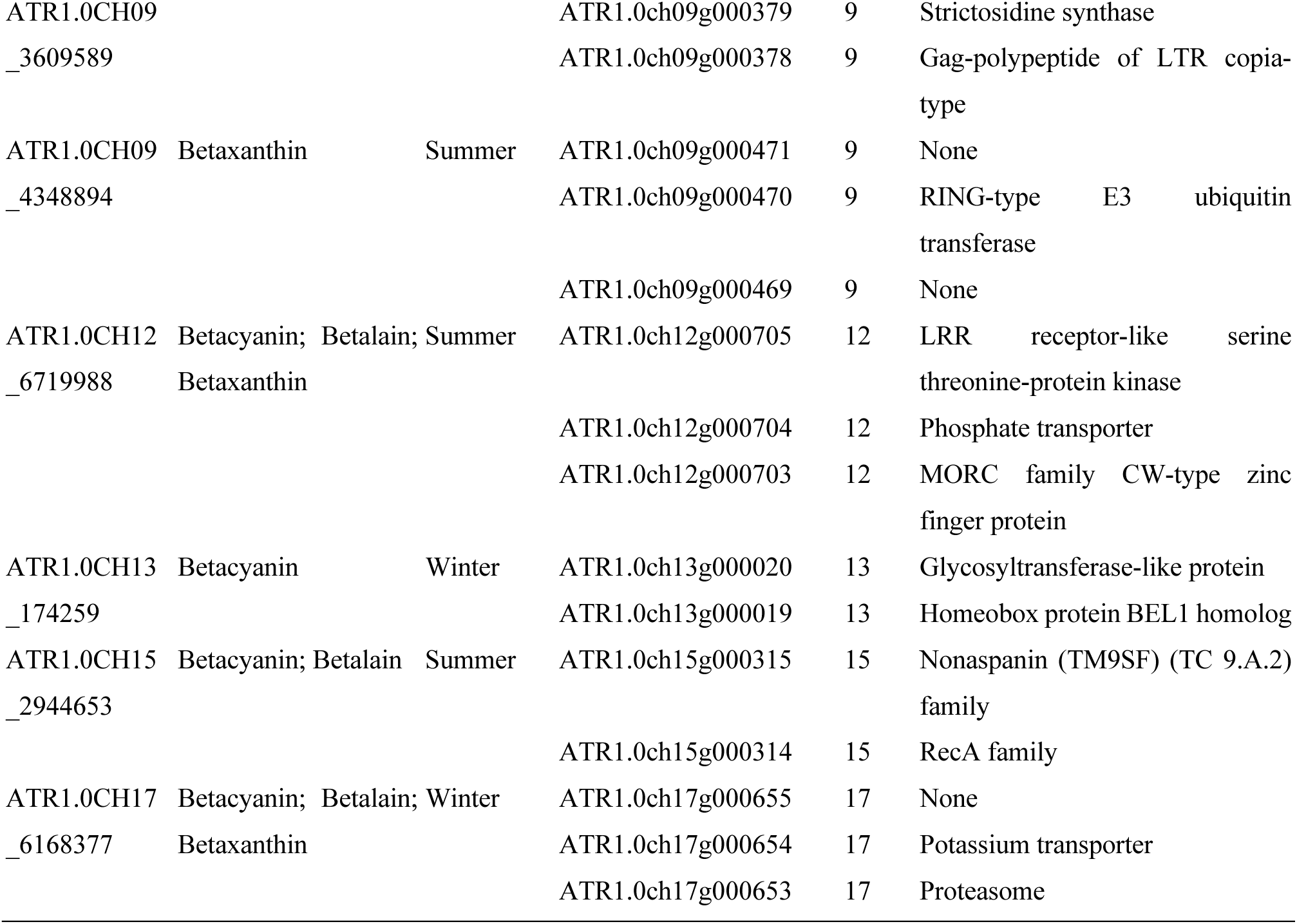
Twelve significant SNPs detected consistently by both FarmCPU and Blink models.

### Differential Gene Expression Under Heat Stress

To investigate the genetic mechanisms underlying betalain biosynthesis under varying temperature conditions, RNA-seq experiments were conducted on Am087 and Am127, two accessions that displayed relatively consistent performance in field trials and higher betalain contents across seasons. The cultivated line Bayam was used as a control. One month after sowing, plants subjected to heat stress exhibited typical stress phenotypes, including stem elongation, delayed panicle production, reduced pigmentation, leaf abscission, senescence, and curling (Figure 4). The decrease in betalain biosynthesis was consistent in Am087 and Am127 but not in Bayam (Supplementary Table 7). This was consistent with the similar changes of betalain content in Am087 and Am127. Overall, 3, 506 and 771 unique differentially expressed genes (DEGs) were identified in Bayam, Am087, and Am127, respectively (Supplementary Table 8). A total of 100 DEGs were shared between Am087 and Am127 (Figure 5A), suggesting that Bayam is more tolerant to heat stress than Am087, which in turn is more resistant than Am127. Both lines exhibited enrichment in genes related to transmembrane transport activity (Supplementary Figure 6), consistent with the expected impact of heat stress on transmembrane stability.

**Figure 4.**
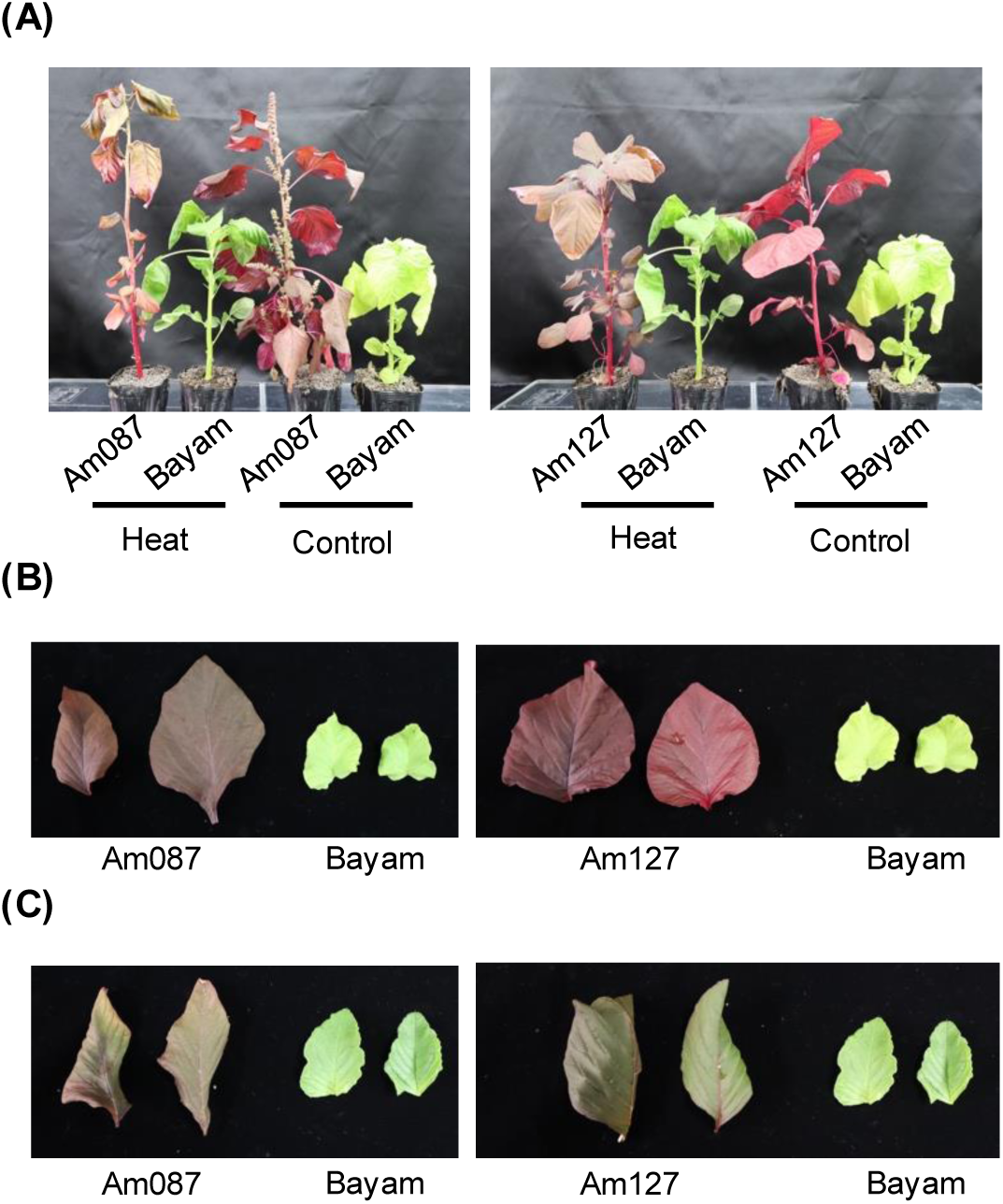
Morphological comparison of Bayam, Am087, and Am127 under control and heat stress conditions. (A) presents the whole plant morphology. (B) and (C) display the leaf morphology under control and heat stress conditions, respectively.

**Figure 5.**
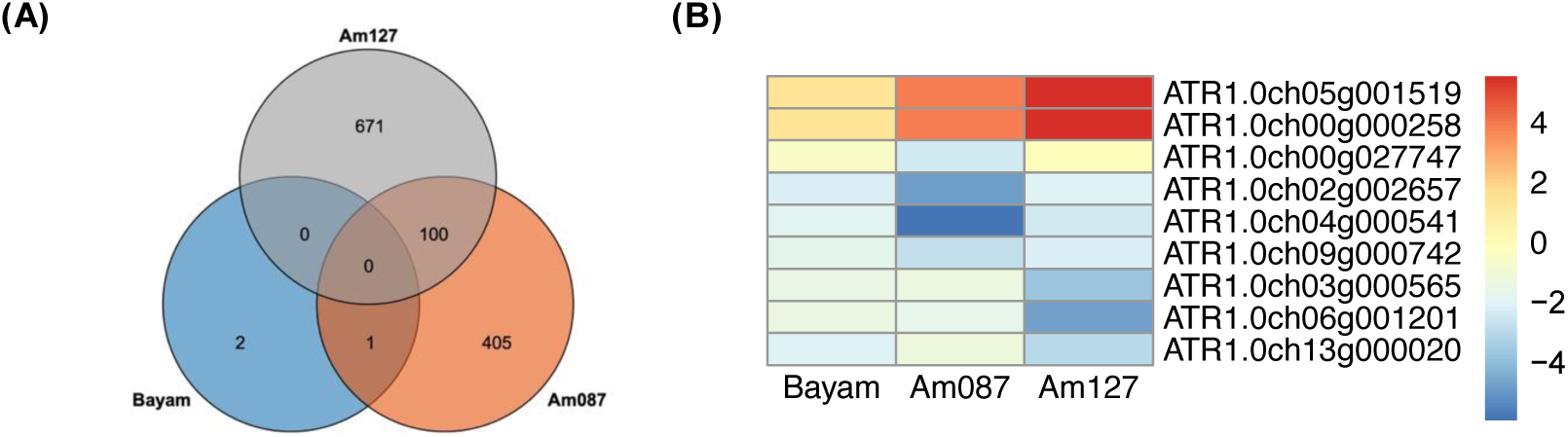
Transcriptomic profiles of the three lines. (A) is the Venn plot of these three lines. (B) is the heatmap of the candidate genes. *ATR1.0ch05g001519* and *ATR1.0ch00g000258* are homologs of *cDOPA*, while *ATR1.0ch00g027747* is a homolog of *CYP76AD1*. Additionally, *ATR1.0ch02g002657*, *ATR1.0ch04g000541*, and *ATR1.0ch09g000742* are homologs of *5GT*. The genes *ATR1.0ch03g000565*, *ATR1.0ch06g001201*, and *ATR1.0ch13g000020* are DEGs associated with significant SNPs under optimal conditions. In the heat map, a redder color indicates higher expression levels under heat stress, while a bluer color indicates lower expression levels under control conditions.

Six homologous genes involved in betalain biosynthesis—two *cDOPA* homologs (*ATR1.0ch05g001519*, *ATR1.0ch00g000258*), one *CYP76AD1* homolog (*ATR1.0ch00g027747*), and three *5GT* homologs (*ATR1.0ch02g002657*, *ATR1.0ch04g000541*, *ATR1.0ch09g000742*)—were differentially expressed in Am087 and Am127 during summer. In Am087, *CYP76AD1* was significantly downregulated, while the *5GT* homologs were also downregulated, despite the upregulation of *cDOPA* homologs (Figure 5B, Table 2). Am127 exhibited similar trends, although without statistical significance (Table 2). These results suggest that under heat stress, the downregulation of *CYP76AD1* and *5GT* homologs leads to reduced betacyanin and betaxanthin levels, with the upregulation of *cDOPA* unable to compensate for this decrease.

**Table 2.**
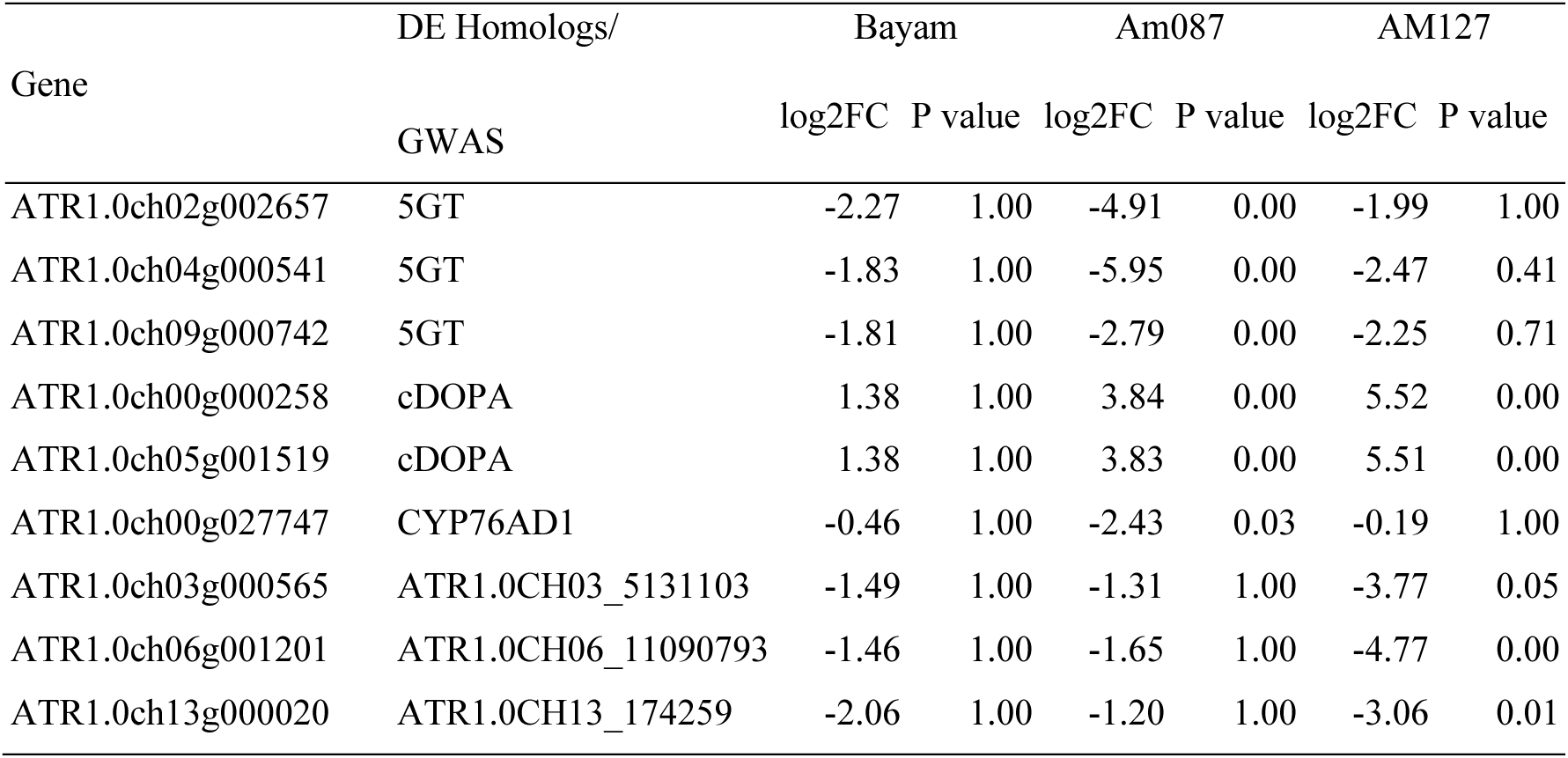
Differential expression of the candidate genes under control and heat stress conditions.

| Gene | DE Homologs/ | Bayam |  | Am087 |  | AM127 |  |
| --- | --- | --- | --- | --- | --- | --- | --- |
|  | GWAS | log2FC | P value | log2FC | P value | log2FC | P value |
| ATR1.0ch02g002657 | 5GT | -2.27 | 1.00 | -4.91 | 0.00 | -1.99 | 1.00 |
| ATR1.0ch04g000541 | 5GT | -1.83 | 1.00 | -5.95 | 0.00 | -2.47 | 0.41 |
| ATR1.0ch09g000742 | 5GT | -1.81 | 1.00 | -2.79 | 0.00 | -2.25 | 0.71 |
| ATR1.0ch00g000258 | cDOPA | 1.38 | 1.00 | 3.84 | 0.00 | 5.52 | 0.00 |
| ATR1.0ch05g001519 | cDOPA | 1.38 | 1.00 | 3.83 | 0.00 | 5.51 | 0.00 |
| ATR1.0ch00g027747 | CYP76AD1 | -0.46 | 1.00 | -2.43 | 0.03 | -0.19 | 1.00 |
| ATR1.0ch03g000565 | ATR1.0CH03_5131103 | -1.49 | 1.00 | -1.31 | 1.00 | -3.77 | 0.05 |
| ATR1.0ch06g001201 | ATR1.0CH06_11090793 | -1.46 | 1.00 | -1.65 | 1.00 | -4.77 | 0.00 |
| ATR1.0ch13g000020 | ATR1.0CH13_174259 | -2.06 | 1.00 | -1.20 | 1.00 | -3.06 | 0.01 |

Considering that our SNP density is approximately one marker per 200 Kb, which is insufficient to adequately cover the gene density of about one gene per 10 Kb, we integrated DEGs to refine the candidate genes identified through GWAS. This approach is based on the rationale that a gene associated with betalain biosynthesis in only one season, and concurrently identified as a DEG, is likely involved in the regulation of betalain synthesis during that specific season. Three significant SNPs identified in the winter betacyanin GWAS were associated with three DEGs (*ATR1.0ch03g000565*, *ATR1.0ch06g001201*, *ATR1.0ch13g000020*), which were significantly downregulated in Am127 under heat stress (Figure 5B, Supplementary Table 8). Given the observed decrease in betacyanin under heat stress, these three genes are implicated in betacyanin biosynthesis and are likely suppressed by elevated temperatures.

## Discussions

### Heat Stress Reduces Betalain Content in Amaranthus tricolor

Heat stress has been observed to induce significant phenotypic abnormalities in several amaranth species during both vegetative and reproductive developmental stages. These abnormalities include reduced leaf pigmentation, decreased plant yield, stunted growth, delayed and deformed panicle emergence, and diminished seed yield (Reyes-Rosales et al., 2023). Despite being one of the few vegetables capable of growing during the summer in tropical regions, there has been limited research on the response of A. tricolor to heat stress. In this study, we found that amaranths produce less betalain in the summer compared to the winter, consistent with previous findings (Reyes-Rosales et al., 2023). The results of this betalain assessment provide valuable insights for breeders aiming to enhance or stabilize betalain production in high-yield lines, thereby contributing to the development of climate-resilient vegetables. Notably, the elite cultivar ’Bayam’ exhibited only three DEGs under varying temperature conditions, with betalain content remaining consistent under both control and heat stress conditions. This suggests that ’Bayam’ is a heat-insensitive line, serving a promising genetic background for the improvement of other traits in *A. tricolor*. Nevertheless, the relatively lower betalain content in Bayam in the winter season still requires improvement.

### The Disruption of Betalain Biosynthesis Under Heat Stress Conditions

Previous studies have established that *CYP76AD1* is crucial for the initial step of betalain biosynthesis, which proceeds either through *DODA* and *5GT* or *CYP76AD1* and *cDOPA* to produce betanin, and via *DODA* to synthesize betaxanthin (Chang et al., 2021). In the current study, we observed that the homologs of *CYP76AD1* were downregulated under heat stress, indicating that betalain biosynthesis is inhibited by heat stress from the earliest stages. Similar downregulation of *CYP76AD1* under high temperatures has been documented in *Suaeda salsa* (Li et al., 2023), suggesting that this may be a common mechanism for the degradation of betalain biosynthesis across species. The biosynthesis of betacyanin via *5GT* also appears to be suppressed under heat stress, as evidenced by the downregulation of *5GT*. Furthermore, the upregulation of *cDOPA* was insufficient to compensate for the downregulation of *5GT*, resulting in lower betacyanin levels under heat stress. Similarly, the reduction in betaxanthin can be attributed primarily to the decreased expression of *CYP76AD1*, as *DODA* expression did not differ significantly between control and heat stress conditions. These findings underscore the critical roles of *CYP76AD1*, *5GT*, and *cDOPA* in regulating betalain biosynthesis. Nevertheless, the betacyanin in Am087 decreases more severely than in Am127, suggesting the downregulation of *CYP76AD1* and *5GT*, may be triggered by other upstream factors.

### Candidate Genes Identified by GWAS

A significant SNP, ATR1.0CH14_4712147, is located within *ATR1.0ch14g000595*, which encodes a DEAD-box ATP-dependent RNA helicase. DEAD-box RNA helicases are crucial for plant development, influencing both vegetative and reproductive growth, as well as mediating responses to abiotic and biotic stress (Li et al., 2023). For example, *OsBIRH1* in rice encodes a DEAD-box RNA helicase that is vital for tolerance to both biotic and abiotic stresses (Li et al., 2008). Similarly, the *Thermotolerant Growth Required 1* in Chinese cabbage has been associated with heat stress tolerance (Yarra and Xue, 2020). In Arabidopsis, T-DNA insertion mutants of DEAD-box RNA helicases show upregulation of *MYB* genes under abiotic stress. *MYB* transcription factors are known to regulate anthocyanin biosynthesis (Petroni and Tonelli, 2011). The *MYB* gene, which directly regulates both *CYP76AD1* and *DODA* in *B. vulgaris* (Hatlestad et al., 2015), is located on Chromosome 2 and is linked to the *CYP76AD1* locus (Goldman and Austin, 2000). In pitaya, *HuMYB132* has been demonstrated to promote betacyanin biosynthesis by binding to *HuCYP76AD1–1* and *HuDODA1* (Xie et al., 2023). In this study, seven *MYB* family genes were significantly differentially expressed in Am127, with six of them downregulated during the summer season. Although *ATR1.0ch14g000595* was not identified as a DEG, genetic variation at this locus may potentially alter its structure, leading to the downregulation of *MYB* genes. Furthermore, earlier research has indicated that *WRKY* genes regulate pigment biosynthesis by interacting with *MYB* genes (Li et al., 2020; Xie et al., 2023). However, in this study, we identified three *WRKY* genes as DEGs, all of which were upregulated under heat stress conditions. This finding contrasts with previous results, suggesting a complex regulatory mechanism governing betalain biosynthesis under abiotic stress.

## Conclusions

This study has demonstrated a notable decrease in betalain content in amaranths during the summer season. Through the integration of genome-wide association studies (GWAS) and differential gene expression (DEG) analysis, we have identified three candidate genes implicated in betalain biosynthesis. Among these, ATR1.0CH03_5131103, which co-localizes with the E3 ubiquitin-protein ligase ORTHRUS, was significantly detected during the winter season, suggesting that ORTHRUS plays a critical role in sustaining betalain biosynthesis under cooler conditions. Additionally, the homologs of *CYP76AD1* and *5GT* were found to be downregulated under heat stress, indicating their responsibility in the reduction of betacyanin and betaxanthin, respectively. Although the homologs of *cDOPA* were upregulated, this upregulation was insufficient to counterbalance the effects of *CYP76AD1* and *5GT* downregulation on betalain content.

### Data availability

Raw sequence reads were deposited in the Sequence Read Archive (SRA) database of the DNA Data Bank of Japan (DDBJ) under the accession number DRA014927, DRA014962, and DRA014963. Assembled sequences are available at DDBJ (accession numbers BSCK01000001 - BSCK01001969) and Plant GARDEN (https://plantgarden.jp).

## Acknowledgments

This work was supported by JSPS KAKENHI (Grant Numbers 19KK0151, 22H05172, and 22H05181) and the Kazusa DNA Research Institute Foundation. We thank the staff in the Flagship of Healthy Diets and Vegetable Diversity and Improvement in the World Vegetable Center and in Kazusa DNA Research Institute for their technical support. We also thank the long-term strategic donors to the World Vegetable Center, including the governments of Taiwan, Germany, Thailand, the Philippines, South Korea, Japan, UK, USAID, and ACIAR.

## Author contributions statement

Y.-P. Lin designed and performed research, analyzed data, and wrote the paper. T. Ohokubo performed research. A. Nashiki performed research and analyzed data. Y. Yoshioka performed research and analyzed data. Y.-Y. Zhang performed research. H.-M. Ting performed research. Sachiko Isobe designed and performed research. Kenta Shirasawa designed and performed research and wrote the paper. Ken Hoshikawa designed and performed research, analyzed data, and wrote the paper.

## Additional Information

Competing financial interests.

The authors declare that they have no competing interests.

## Legends of supplementary data

**Supplementary Figure 1 Comparisons of betalain between seasons and leaf colors.**

**Supplementary Figure 2 The genome size of Bayam.** K-mer analysis indicates the genome size of Bayam is about 596.4 Mb

**Supplementary Figure 3 Genome duplication in the *A. tricolor* cv. Bayam.** Similar genome segments are connected with lines.

**Supplementary Figure 4 Principal Coordinate Analysis of the core collection.**

**Supplementary Figure 5 Manhattan plots of betaxanthin (A480), betacyanin (A538), and betalain** A) The Manhattan plots of the FarmCPU model. B) The Manhattan plots of the Blink model.

**Supplementary Figure 6 GO enrichment analyses.** A) and B) are the top 30 enriched GO in Am087 and Am127, respectively. C) is the enriched GO shared by Am087 and Am127.

**Supplementary Table 1 Phenotypes of the core collection under the two seasons.**

**Supplementary Table 2 Statistics of the genome assembly of *A. tricolor* cv. Bayam.**

**Supplementary Table 3 Statistical assembly summary of each chromosome**

**Supplementary Table 4 The homologous genes of the five enzymes in the *A. tricolor* genome.**

**Supplementary Table 5 Significant SNPs of the betalain contents in the two seasons detected by FarmCPU and Blink.**

**Supplementary Table 6 Candidate genes directly targeted by or adjacent to the significant SNPs.**

**Supplementary Table 7 Betacyanin and betaxanthin of *A. tricolor* cv. Bayam, Am087, and Am127 under the control and heat conditions.**

**Supplementary Table 8 DEGs in Bayam, Am087, and Am127 and their annotations.**

